# Sex differences in the effects of enhanced adult hippocampal neurogenesis on pattern discrimination and Intellicage spatial learning in Bax-knockout mice

**DOI:** 10.64898/2026.11.23.768231

**Authors:** Arasely M. Rodriguez, Elif Tunc-Ozcan

**Author notes:** Corresponding Author: Elif Tunc-Ozcan, Dept. Neurosciences, MSC08 4740, 1 University of New Mexico, Albuquerque, NM 87131-0001. Funding Sources: NIMH 5R00MH125016, NIAAA 1F31AA031920.

## Abstract

Adult hippocampal neurogenesis has been implicated in learning and memory. Although increasing neurogenesis has been associated with improved spatial learning and pattern discrimination, whether these benefits differ between male and female mice remains unclear. The goal of this study was to see if increasing adult hippocampal neurogenesis improves learning in male and female mice, and whether the benefit is the same in both sexes. We used conditional deletion of the pro-apoptotic gene Bax to increase the survival of newborn dentate granule cells. Male and female BAX knockout (KO) and wild-type (WT) mice were tested in Intellicage place learning and reversal learning tasks, and an A/B context discrimination task. BAX KO mice had more tdTomato-positive newborn dentate granule cells than BAX WT mice, with no significant effect of sex. During Intellicage adaptation, there were no differences between genotypes or sexes in total visits and licks. During place learning, male mice had a higher percent correct response than female mice, while female mice showed increases in licks and correct visits from day 1 to day 4. BAX KO mice made more visits overall but had a lower percent correct response than BAX WT mice on day 4. During reversal learning, male BAX KO mice showed a decrease in visits and had fewer visits with a lick than male BAX WT mice on day 4. In the A/B task, male BAX KO mice had higher discrimination scores than male BAX WT mice on day 5, while female BAX KO mice showed an initial advantage over female BAX WT mice. These results partially support our hypothesis that increasing neurogenesis would benefit male mice more. Overall, increasing the survival of newborn dentate granule cells does not improve learning in the same way across sexes and tasks.

## Introduction

Mouse behavioral tasks are widely used in preclinical studies to assess the effect of genetic alterations. However, behavioral performance can be influenced by factors beyond experimental manipulation, potentially complicating interpretation of the behavioral test results, and reducing reproducibility. For example, familiarity with the testing environment can affect behavioral outcomes, particularly in tasks assessing stress- and anxiety-related behaviors (Kas and Van Ree, 2004; Kyriakou *et al*., 2018). Interaction between the rodent and experimenter has also been shown to alter behavior results (Sousa, Almeida and Wotjak, 2006). Automated home-cage testing offers an approach to reducing some of these influences by assessing behavior in a familiar environment while limiting experimenter involvement (Grieco *et al*., 2021).

The Intellicage system is a home-cage testing system that enables behavioral assessment of group-housed mice. In the Intellicage system, there are four operant corners that have an RFID sensor at their entrance. Inside each corner there are two water sipper ports that are gated. At each gate there is a touch sensor that can be activated by a mouse’s nose. This system is inside a standard large rodent cage; it allows for the uninterrupted recording of these animals across multiple days. The Intellicage can be used to assess spatial learning through tasks in which mice learn to obtain water at designated locations (Kiryk *et al*., 2020). Meta-analyses of conventional spatial tasks have reported an overall male advantage in rodents (Jonasson, 2005).

Such task dependent differences highlight the importance of considering sex as a biological variable. Despite its potential influence on behavior, sex has historically been underexamined in preclinical research. Male-only study designs were often justified by concerns that fluctuating ovarian hormones would increase variability in females (Weigard, Loviska and Beltz, 2021). However, excluding females limits generalizability of findings and prevents the identification of sex-dependent responses. Sex can influence disease processes and responses to therapeutic interventions, as well as behavioral performance (Hamson, Roes and Galea, 2016). Including both males and females is therefore important for determining whether manipulations targeting learning and memory have comparable effects across sexes.

Specific types of hippocampal learning can be altered by adult hippocampal neurogenesis. Adult hippocampal neurogenesis is the generation and integration of new neurons in the adult dentate gyrus. Evidence that increasing neurogenesis can improve performance in some learning tasks comes from studies using genetic manipulations. For example, Barth et al. (2023) reported that deletion of epidermal growth factor-like protein 7 (EGFL7) using a Nestin-CreERT2 model increased adult hippocampal neurogenesis (AHN) and improved performance in both the Morris water maze and an Intellicage place learning task; they only looked at male mice. Another approach involved the conditional deletion of pro-apoptotic gene Bax. Like the knockout of EGFL7, Bax knockout also increases AHN. Studies using this approach have reported increased neurogenesis and improvements in pattern discrimination learning (Sahay *et al*., 2011; used both sexes but did not look at differences; Besnard and Sahay, 2021; only used male mice). Pattern discrimination learning involves the ability to distinguish between two similar but different contexts. However, whether the behavioral effect of increased neurogenesis differ between males and females remains insufficiently understood.

Together, evidence for task-dependent sex differences in spatial learning and behavioral effects of increased neurogenesis motivates examining these factors jointly. Therefore, the present study investigated whether increasing adult hippocampal neurogenesis through Bax knockout (KO) differentially affects place learning and pattern separation in male and female mice. We hypothesize that increasing neurogenesis will produce greater performance in these tasks in male mice than females.

## Methods

### Animals

Adult (PND 56-125) Transgenic NestinCreERt2:tdTomato:BAXf/f (KO) and Nestin-Cre^T2^:tdTomato:BAXwt (WT) mice on a C57BL/6J background, female and males, were used in these studies. Counts were: Female BAX KO = 11, Female BAX WT = 8; Male BAX KO 7; Male BAX WT = 12. These mice strains were isolated from the same founder crosses. Nestin-Cre^T2^:tdTomato mice were previously described (Gustus *et al*., 2020; Rodriguez *et al*., 2026). BAXf/f mice were generously provided by Dr. Amar Sahay (Sahay et al., 2011; Besnard & Sahay 2021). NestinCreERt2:tdTomato mouse strain expresses tdTomato in the progeny of Nestin+ cells, thus labeling newborn dentate granule cells. When BAXf/f is present, these labeled cells also have the BAX protein knocked out. Thus, in Nestin-Cre^T2^:tdTomato:BAXf/f newborn dentate granule cells can be quantified and also the number of these cells increases compared to the BAX WT line as previously described by Sahay et al. (2011). Genotyping was routinely performed on tail tissue by qualitative PCR services performed by TransnetYX genotyping. Mice were housed in a 12:12 normal light cycles with lights off at 18:00 and in a humidity and temperature-controlled room. Mice were genotype-segregated, sex-segregated and group housed. Mice had ad-libitum access to standard mice chow and unless in the Intellicage they had ad-libitum access to water. All protocols were approved by the Institutional Animal Care and Use Committee of the University of New Mexico and conformed to the National Institutes of Health Guide for the Care and Use of Laboratory Animals.

### Tamoxifen

Tamoxifen (Sigma-Aldrich, St. Louis, MO, USA) was dissolved in sunflower oil (Sigma-Aldrich; 30 mg/mL) via a heated water bath sonicator. All mice received tamoxifen injection (i.p) for 5 consecutive days at a dose of 180 mg/kg.

### Intellicage

2 weeks prior to placement in Intellicage mice were subcutaneously implanted with RFID chips in the dosro-cervical region. The IntelliCage apparatus (TSE Systems; Berlin, Germany) is a 39 cm x 58 cm x 21 cm cage that contain four operant corners. These corners have two water stations that can be accessed through the lift of a gate, at the side of each of these gates is a touch sensor. The corners also have an RFID sensor at every corner entrance that allows for the tracking of mice via the implanted RFID. There are no barriers to enter a corner; the only gates are located at water-access ports.

### Free Adaptation (FA)

Eight weeks after tamoxifen administration mice were placed in the Intellicage. During the first day of placement into the Intellicage all water gates were set to open allowing mice to acclimate the environment. This was done for a single day.

### Door Adaptation (DA)

All water access gates were closed. Registration of a mouse in a corner would open the door. This was done to acclimate mice to the gates. There were no door specific assignments. This was done for a single day.

### Nosepoke Adaptation (NPA)

All water access gates were closed. A nosepoke at the touch-sensor of a cage would open the gate. Water access was limited to dark cycle (1800-0600). This was so mice were trained to access water after performing an action and only during a specific time period. Any mouse that did not successfully lick water during these past three days was removed from the experiment. Gates only open when one mouse is registered in a corner to prevent access by non-nosepoking mice. There were no specific door assignments. This was done for a single day.

### Place Learning (PL) and Reversal Learning (RL)

All water access gates were closed. The least visited corner during the last three days by the mouse was assigned as the water access corner, the correct corner. Water access was limited to dark cycles at the assigned door. At these correct corners a nosepoke was needed to open the water gate; a nosepoke in an incorrect corner would not open the water gate. Selected corners were balanced across mice so as to limit imitation learning and overcrowding. Place learning was conducted for four days. After which, the corner opposite to the corner during place learning was designated as the new correct corner to assess the reversal learning. Reversal learning was conducted for four days.

### Non-Intellicage Behavior Testing

1 week prior to Intellicage placement mice were administered an A/B context discrimination task. In this task there are two boxes: (A) shock box with thick and thin alternating bar floor (B) an identical box but with a flat floor, semi-circular boundaries, and striped walls. Prior to A/B, mice were acclimated to behavioral room for 1 hr. Mice were placed in box A where they received a 2s (0.5 mA) shock after 90 seconds, they received a second shock after another 90 seconds, mice remained for 30 seconds after the last shock. Mice were returned to their home cage. The boxes were disinfected with 70% ethanol. After 1 hr, mice were placed in box B, where they did not receive any shock and remained in the box for 210 seconds. This procedure was followed for 5 days, alternating the initial box. Mice received a shock every day in the shock box.

After the last day of the Intellicage reversal learning, mice were run through an open-field test and a tail-suspension test. One hour prior to these tests mice were acclimated to the behavior room. The OFT was conducted in a white square arena (30 cm x 30 cm). Light was adjusted so that the corners of the arena were ∼45 lux and the center of the box was ∼90 lux. The box was disinfected with 70% ethanol. Mice were released into the center of the arena and recorded for a total of 8 min. The same day mice were run in a TST. Mice were hung onto a bar with tape. A cone was placed over the tail to prevent tail climbing. Mice were assessed for a duration of 5 min. Quantification of behavior was conducted using video-tracking software, Ethovision 18 XT (Noldus, Leesburg, VA).

### Neurogenesis Quantification

Mice were euthanized via CO2 inhalation. They were perfused transcardially with 0.1% procaine (Sigma-Aldrich and 2 U/mL of heparin (Fresenius Kabi USA, Lake Zurich, IL) in phosphate buffered saline (PBS), then with a 4% paraformaldehyde (PFA) solution in PBS. Brains were kept in 4% PFA for 24 hrs. and subsequently placed in a 30% sucrose in PBS solution until they sunk. Brains were coronally sectioned at 40 um at the dorsal hippocampus. Sections were mounted with DAPI Flouromount G (Electron Microscopy Sciences, Hatfiel, PA). Images were taken at 20x on a Zeiss LSM800 confocal microscope. tdTomato positive cells within the granular zone were manually counted while blinded using QuPath v0.5.1 (Bankhead *et al*., 2017). The average number of tdTomato positive cells was taken across three sections.

### Statistical Analysis

We calculated Intellicage data following Judd et al (2024) and is as follows:

Total Visits = total number of visits to all corners.

Total Licks = total number of licks made. Measure of water consumption.

Total Correct Visits = total number of correct corner visits.

Total Visits with Lick = total number of visits where at least one lick occurred. This indicates that mice performed the correct actions to receive water.

Percent Correct Response = Number of visits with (>=) 1 lick (i.e. correct response) / over total visits

For other behaviors:

TST –percent immobile =time immobile / total time in TST

OFT – percent distance in center = distance in center / total distance in center (cm); percent time in center = time in center / total time spent in center (sec.)

A/B – discrimination ratio = (time freezing in context A – time freezing in context B) / (time freezing in context A + time freezing in context B); positive number indicates correct freezing in shock context and negative number indicates freezing in incorrect context (non-shock). Only the time before the first shock was quantified, the first 90 seconds.

Mice that did not make any licks during the adaptation phase were removed from the experiment. Data were analyzed using IBM SPSS (v.32). Generalized linear mixed models were used when there were repeated measures. For single day value for the Intellicage (FA, DA, NPA) a generalized linear model was used; for the OFT, TST, and Neurogenesis a linear model was used. Mean estimated marginal mean (EMM) differences are reported with 95% CI. Pairwise comparisons of EMM with Bonferroni corrections were conducted. Not all pairwise comparisons were graphed.

## Results

### BAX KO mice had increased neurogenesis compared to BAX WT mice

Only a significant effect of genotype was observed on the number of tdTomato+ nDGCs [F (1, 37) = 17.425; p < 0.001] (Fig. 1). BAX KO mice had an average of 36.14 more tdTomato+ nDGCs than BAX WT mice (95% CI [18.543, 53.729]; p < 0.001), indicating enhanced neurogenesis in the BAX KO genotype.

**Figure 1.**
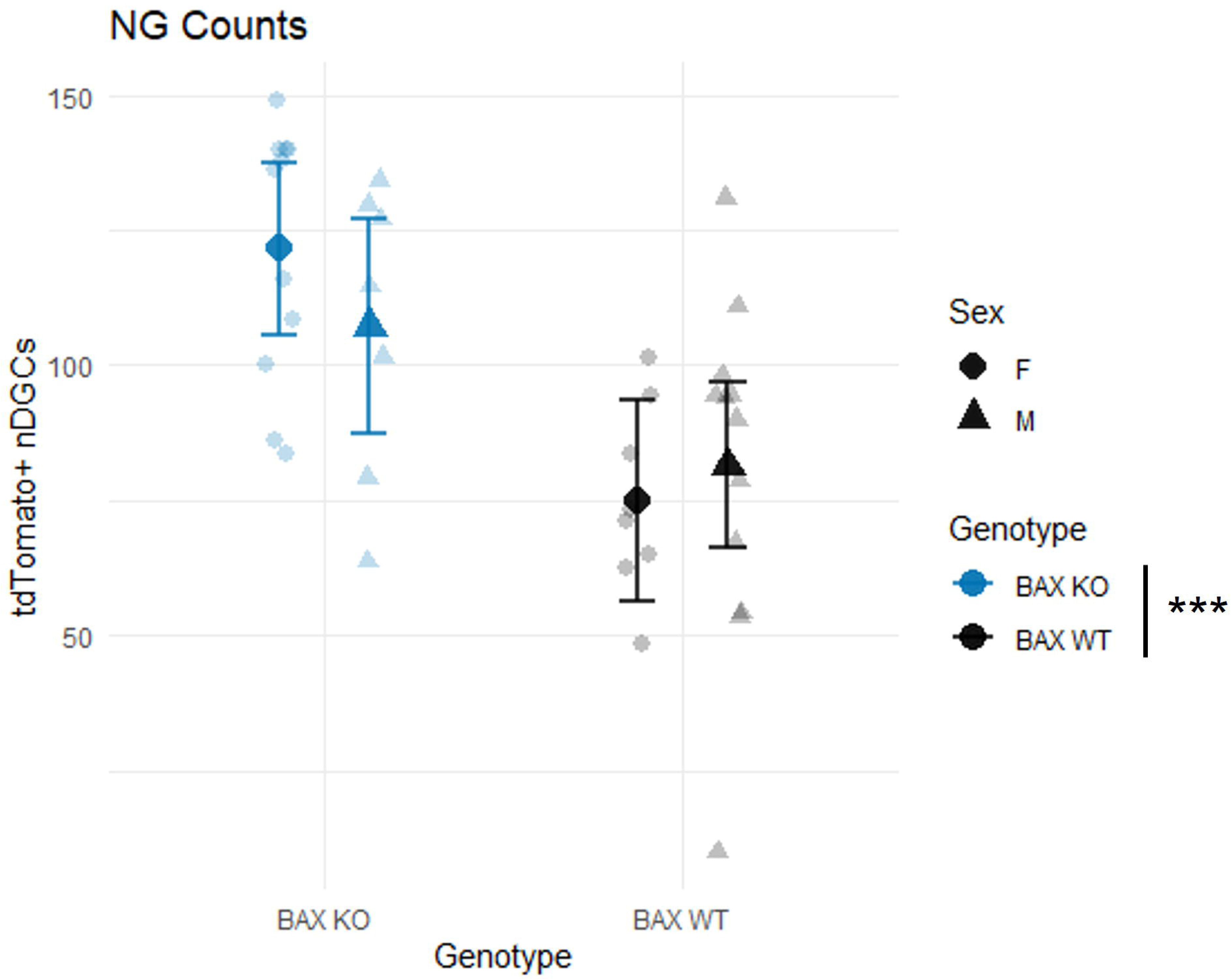
Adult neurogenesis was increased in BAX KO mice compared to BAX WT mice. Number of tdTomato+ newborn dentate granule cells (nDGCs) in the dentate gyrus. Small symbols show individual mice; large symbols show estimated marginal means (EMMs) with 95% confidence intervals (CIs). Circles: females; triangles: males. BAX KO: blue; BAX WT: black. ***p < 0.001, main effect of genotype.

### BAX KO mice and BAX WT mice, regardless of sex, did not differ in the number of visits and licks during Intellicage adaptation

During the free adaptation phase in the Intellicage, there was no significant effect of genotype on total visits (Wald chi-square = 0.06; df = 1; p = 0.812; Fig. 2A), nor was there an effect of sex (Wald chi-square = 1.48; df = 1; p = 0.224). Additionally, there was no interaction between genotype and sex (Wald chi-square = 0.16; df = 1; p = 0.691). Likewise, there were no significant effects of genotype (Wald chi-square = 0.01; df = 1; p = 0.918) or sex (Wald chi-square = 0.69; df = 1; p = 0.403) on the number of licks (Fig. 2B), nor was there an interaction effect (Wald chi-square = 0.43; df = 1; p = 0.512).

**Figure 2.**
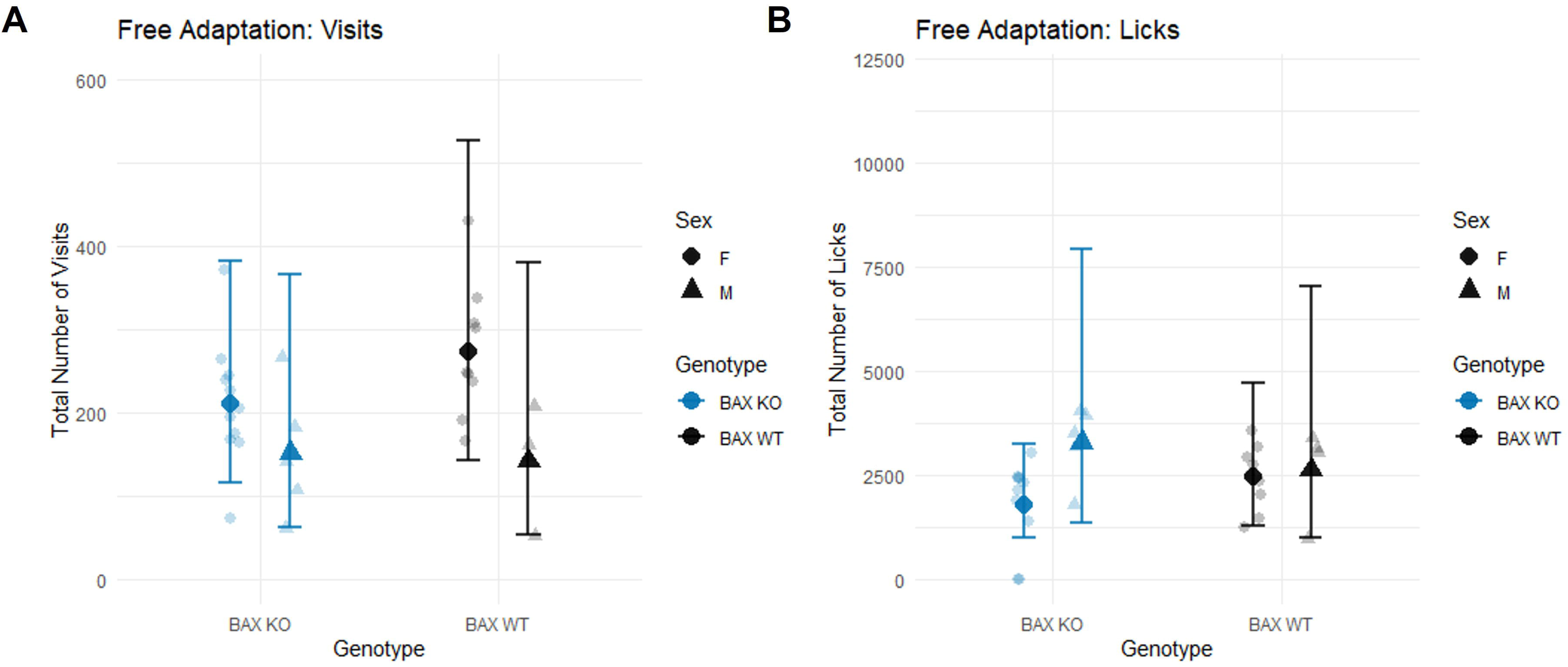
Visits and licks did not differ by genotype or sex during free adaptation. (A) Total number of corner visits and (B) total number of licks during the free adaptation phase in the Intellicage. Small symbols show individual mice; large symbols show estimated marginal means (EMMs) with 95% confidence intervals (CIs). Circles: females; triangles: males. BAX KO: blue; BAX WT: black.

In the door adaptation phase, results showed no significant effect of genotype on total visits (Wald chi-square = 0.54; df = 1; p = 0.463), nor of sex (Wald chi-square = 0.28; df = 1; p = 0.598; Fig. 3A). There was also no interaction between genotype and sex (Wald chi-square = 0.02; df = 1; p = 0.902). Similarly, there were no significant effects on total licks (Fig. 3B) for either genotype (Wald chi-square = 0.01; df = 1; p = 0.906) or sex (Wald chi-square = 0.43; df = 1; p = 0.510), and no interaction effect (Wald chi-square = 0.41; df = 1; p = 0.520).

**Figure 3.**
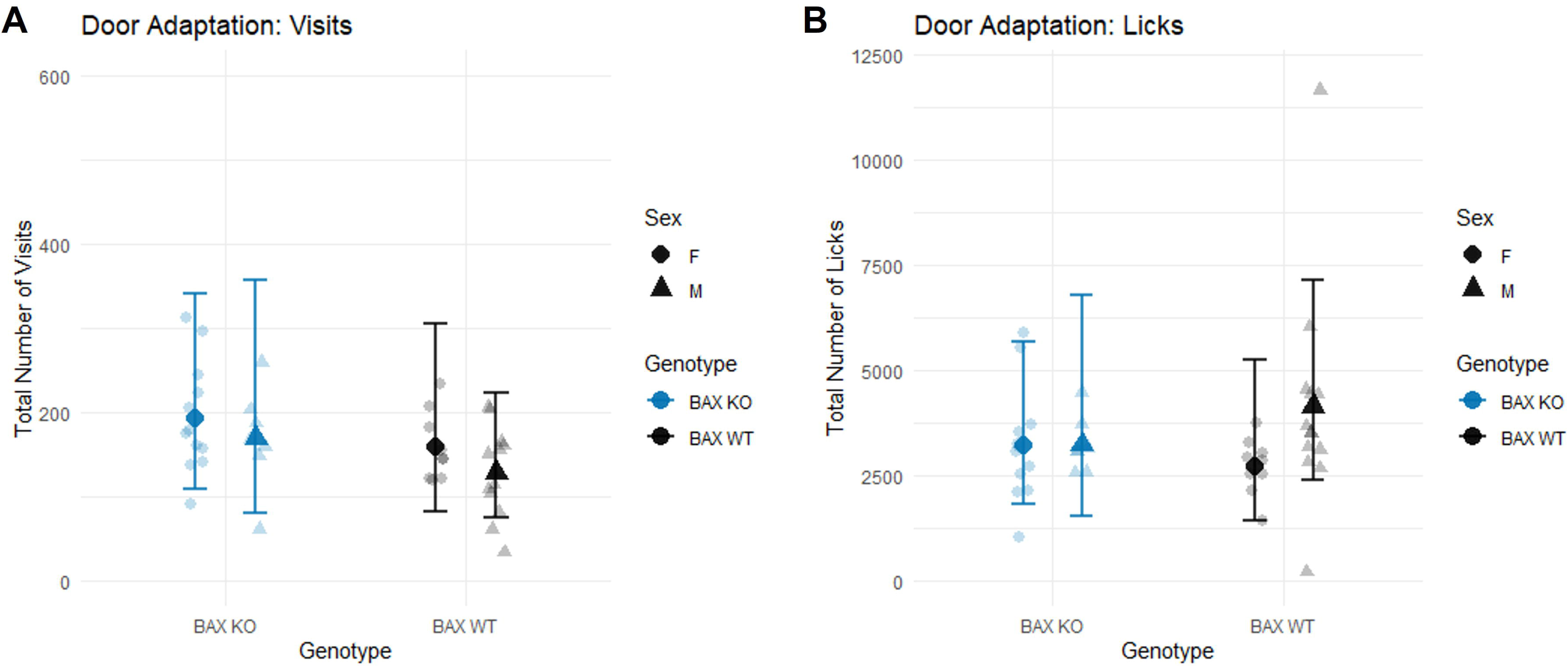
Visits and licks did not differ by genotype or sex during door adaptation. (A) Total number of corner visits and (B) total number of licks during the door adaptation phase in the Intellicage. Small symbols show individual mice; large symbols show estimated marginal means (EMMs) with 95% confidence intervals (CIs). Circles: females; triangles: males. BAX KO: blue; BAX WT: black.

During the nosepoke adaptation phase, there was no significant effect of genotype on total visits (Wald chi-square = 0.42; df = 1; p = 0.517), nor of sex (Wald chi-square = 0.05; df = 1; p = 0.822), with no interaction between genotype and sex (Wald chi-square = 0.12; df = 1; p = 0.731; Fig. 4A). Likewise, there were no effects of genotype (Wald chi-square = 0.08; df = 1; p = 0.777) or sex (Wald chi-square = 0.80; df = 1; p = 0.370) on total licks (Fig. 4B), and no interaction (Wald chi-square = 0.36; df = 1; p = 0.548). Overall, these results indicate that during the adaptation phase BAX KO mice and BAX WT mice did not perform differently. Likewise female and male mice performed similarly.

**Figure 4.**
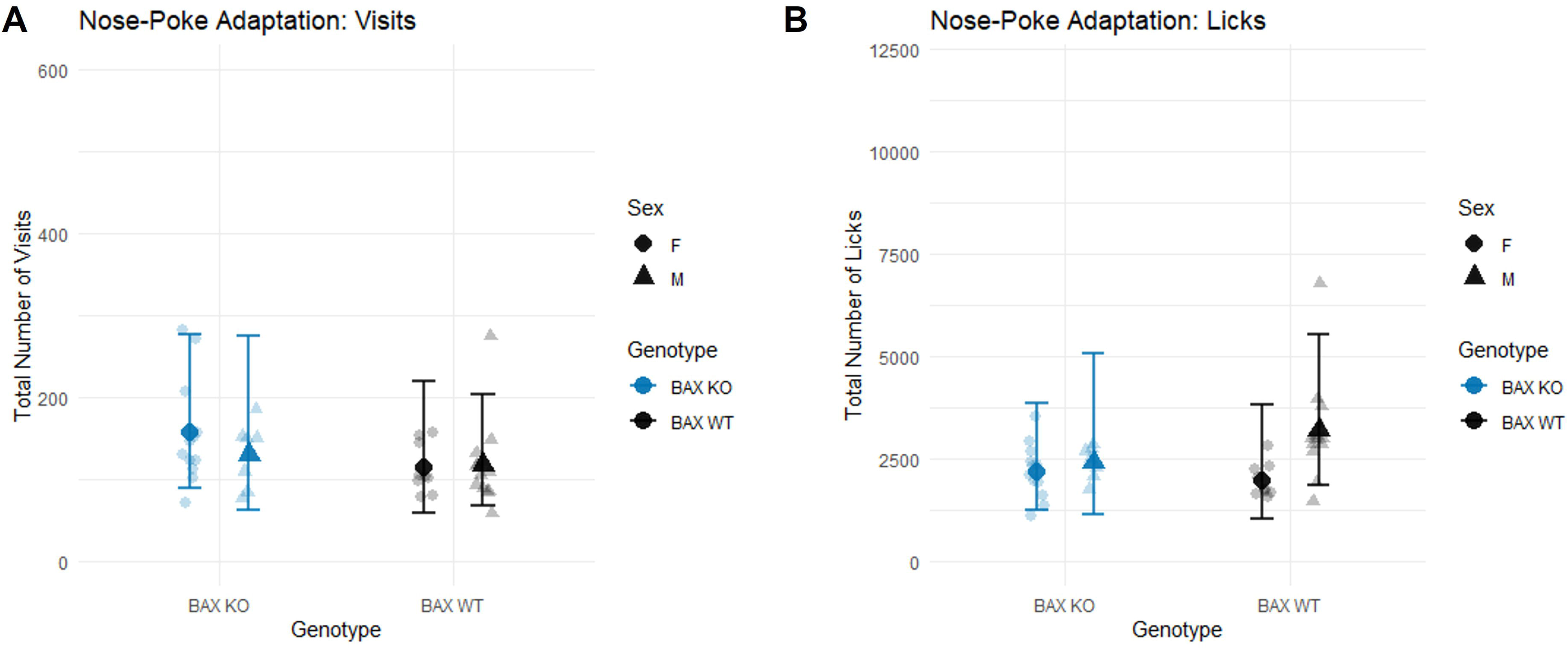
Visits and licks did not differ by genotype or sex during nosepoke adaptation. (A) Total number of corner visits and (B) total number of licks during the nosepoke adaptation phase in the IntelliCage. Small symbols show individual mice; large symbols show estimated marginal means (EMMs) with 95% confidence intervals (CIs). Circles: females; triangles: males. BAX KO: blue; BAX WT: black.

### During place learning all mice developed a preference for the correct corner. Female mice had the most room of improvement due to initial lower performance, however male mice made more percent correct responses

A significant interaction was noted among genotype, sex, and day for total visits [F(3, 46) = 3.534; p = 0.022] (Fig. 5A). Female BAX KO mice had 51.150 more visits than female BAX WT mice on day 4 (95% CI: [8.528, 93.771]; p = 0.019). On day 4, female KO mice made 39.81 more visits than day 1 (95% CI: [3.328, 76.295]; p = 0.025). A significant main effect of genotype was observed [F(1, 37) = 4.532; p = 0.040], with BAX KO mice averaging 24.457 more visits than BAX WT mice across sex and day (95% CI: [0.826, 48.089]; p = 0.043). BAX KO increased the number of visits that mice were making compared to BAX WT mice, this was especially true for female BAX KO mice.

**Figure 5.**
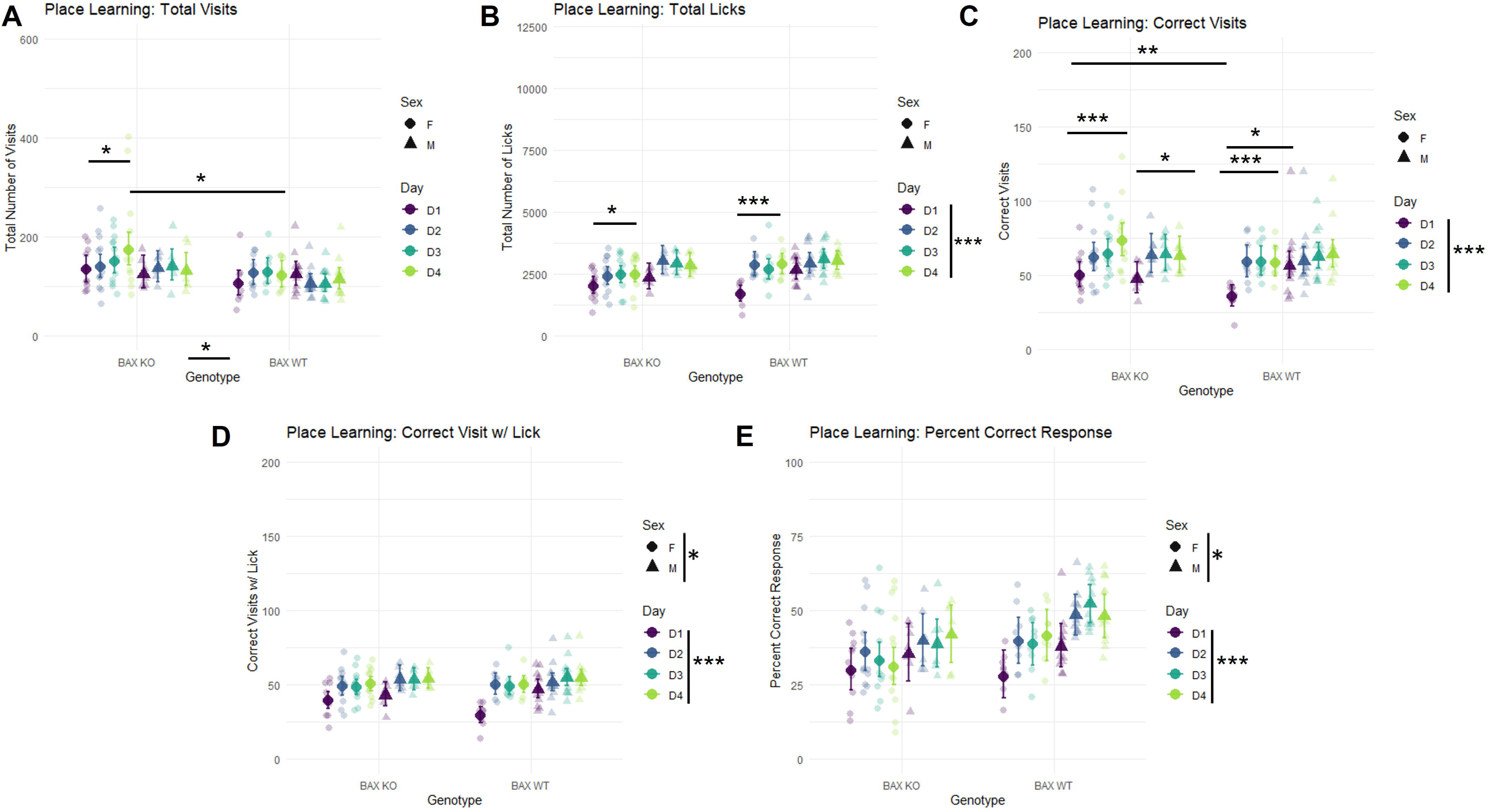
Male mice performed better than female mice in place learning, but female mice improved more across days. (A) Total visits, (B) total licks, (C) correct visits, (D) correct visits with a lick, and (E) percentage of correct responses on days 1–4 (D1–D4). Small symbols, individual mice; large symbols, estimated marginal means (EMMs) with 95% confidence intervals (CIs). Circles, females; triangles, males. Vertical bars next to the keys mark main effects; horizontal bars mark pairwise comparisons. *p < 0.05, **p < 0.01, ***p < 0.001.

A significant interaction of genotype, sex, and day was also significant for total licks [F(3, 55) = 3.273; p =0.028] (Fig. 5B). On day 4, female BAX KO mice made 465.644 more licks than day 1 (95% CI [71.779, 859.510]; p = 0.013). On day 4, female BAX WT mice made 1195.067 more licks than day 1 (95% CI [739.675, 1650.458]; p < 0.001). An interaction of sex and day was observed [F(3, 55) = 2.836; p = 0.046], with male mice showing 659.113 more licks than female mice on day 1 (95% CI: [238.847, 1079.379]; p = 0.003) and 429.503 more on day 3 (95% CI: [16.621, 842.386]; p = 0.042). There was a significant effect of day on the total licks [F(3, 55) = 17.390; p < 0.001], with an increase of 653.466 licks from day 1 to day 4 (95% CI: [414.513, 892.379]; p < 0.001). All mice were making more licks on day 4 than day 1; however male mice made more licks than female mice on day 1. However, the greatest increase in licks were only from day 1 to day 4 were seen in the female BAX KO mice and female BAX WT mice which is probably due to the initial lower licks than male mice.

Furthermore, a significant interaction of genotype, sex, and day were observed for total correct visits [F(3, 46) = 4.489; p = 0.008] (Fig. 5C). On day 1, female BAX KO mice had 14.364 more correct visits than female BAX WT mice (95% CI: [3.620, 25.108]; p = 0.010). Male BAX WT mice made 20.596 more correct visits than female BAX WT mice on the same day (95% CI: [9.538, 31.654]; p < 0.001). On day 4, female BAX KO mice made 23.011 more correct visits than day 1 (95% CI: [11.431, 34.59]; p < 0.001); male BAX KO mice made 15.525 more correct visits (95% CI: [2.387, 28.663]; p = 0.013); female BAX WT mice made 22.857 more correct visits (95% CI: [12.215, 33.067]; p < 0.001); male BAX WT did not have a significant difference D4-D1. Day effects were also significant [F(3, 46) = 27.579; p < 0.001], with mice making 17.659 more correct visits on day 4 compared to day 1 (95% CI: [11.900, 23.419]; p < 0.001). All mice were making more correct visits by day 4. On the first day, female BAX KO mice made more correct visits than female BAX WT mice. However, this difference was attenuated by day 4. Overall, mice made more correct visits by day 4 indicating that they were developing a preference for the correct corner. Interestingly only male BAX WT mice did not make a significant increase in correct visits from day 1 to day 4. However, this was probably due to WT mice reaching a ceiling on day 1, as male BAX WT mice made significantly more correct visits than female BAX WT who did make significant day 1 to day 4 difference.

There was no significant interaction of genotype, sex, day on total correct visits with lick (Fig. 5D). The analysis of total correct visits with a lick revealed a significant interaction among sex and day [F(3, 49) = 3.000; p = 0.039]. On day 1, male mice had 10.940 more correct visits with a lick compared to female mice (95% CI: [4.403, 17.477]; p = 0.001). But there was no difference in day 4. There was also a significant main effect of sex on correct visits with a lick [F(1, 42) = 5.756; p = 0.021]. On average, male mice made 6.297 more correct visits with a lick than female mice (95% CI: [0.966, 11.628]; p = 0.022). A significant main effect of day was also observed [F(3, 49) = 23.560; p < 0.001], with an increase of 13.166 correct visits with a lick on day 4 compared to day 1 (95% CI: [9.178, 17.154]; p < 0.001). Male mice making more correct visits with a lick is in line with male mice making more licks. The initial day and sex effect of male mice making more correct visits with a lick was also probably present due to the lower licks that female mice made on day 1 that allowed for the significant day 1 to day 4 comparison in total licks. Overall, these data support that male mice are performing better than female mice.

There was no significant interaction of genotype, sex, day on percent correct (Fig. 5E). There was a significant interaction between genotype and day for percent correct response [F(3, 51) = 3.171; p = 0.032]. Averaged across sex, BAX KO mice had a decrease of 8.5 percentage points in percent correct response on day 4 compared to BAX WT mice (95% CI: [-0.167, -0.004]; p = 0.040). Additionally, sex had a significant effect on percent correct response [F(1, 37) = 5.762; p = 0.022]. Male mice average 8.2 percentage points higher in correct responses than female mice across genotype and day (95% CI: [0.013, 0.152]; p = 0.021). The main effect of day was significant [F(3, 51) = 9.614; p < 0.001], indicating that day 4 had 7.8 percentage points more correct responses when compared to day 1 (95% CI: [0.031, 0.125]; p < 0.001). Overall, all mice were making more correct visits with a lick over their total visits by day 4, this indicates that they learned which was the correct corner as they had a higher percent correct response. However, interestingly BAX KO mice showed a decrease. But this decrease is likely attributed to the fact that BAX KO mice were making overall more visit corners than it being that BAX KO performed more poorly than BAX WT mice. Looking at all the values, as males made more licks, correct visits, correct visits with a lick, and had a higher percentage of correct responses, male mice learned better than female mice. However, as female BAX KO mice showed the more instances of significant differences from day 1 to day 4, this means that BAX KO helped female mice improve more so than male mice.

### During reversal learning all mice learned the correct corner as they made an increase of licks by day 3, however only BAX WT mice had a higher percent correct response

There was a significant effect of genotype, sex, and day on total visits [F (3, 44) = 3.088; p = 0.037] (Fig. 6A). On day 4, male BAX KO mice made 47.930 fewer visits than day 1 (95% CI: [- 82.344, -13.516]; p = 0.002). A significant effect of day was observed on total visits [F(3, 44) = 7.651; p < 0.001]. The average marginal mean difference on day 4 was 16.804 fewer visits compared to day 1 (95% CI: [-28.557, -5.051]; p < 0.002). These results demonstrate that there is an overall decrease in visits that especially noticeable in male BAX KO mice.

**Figure 6.**
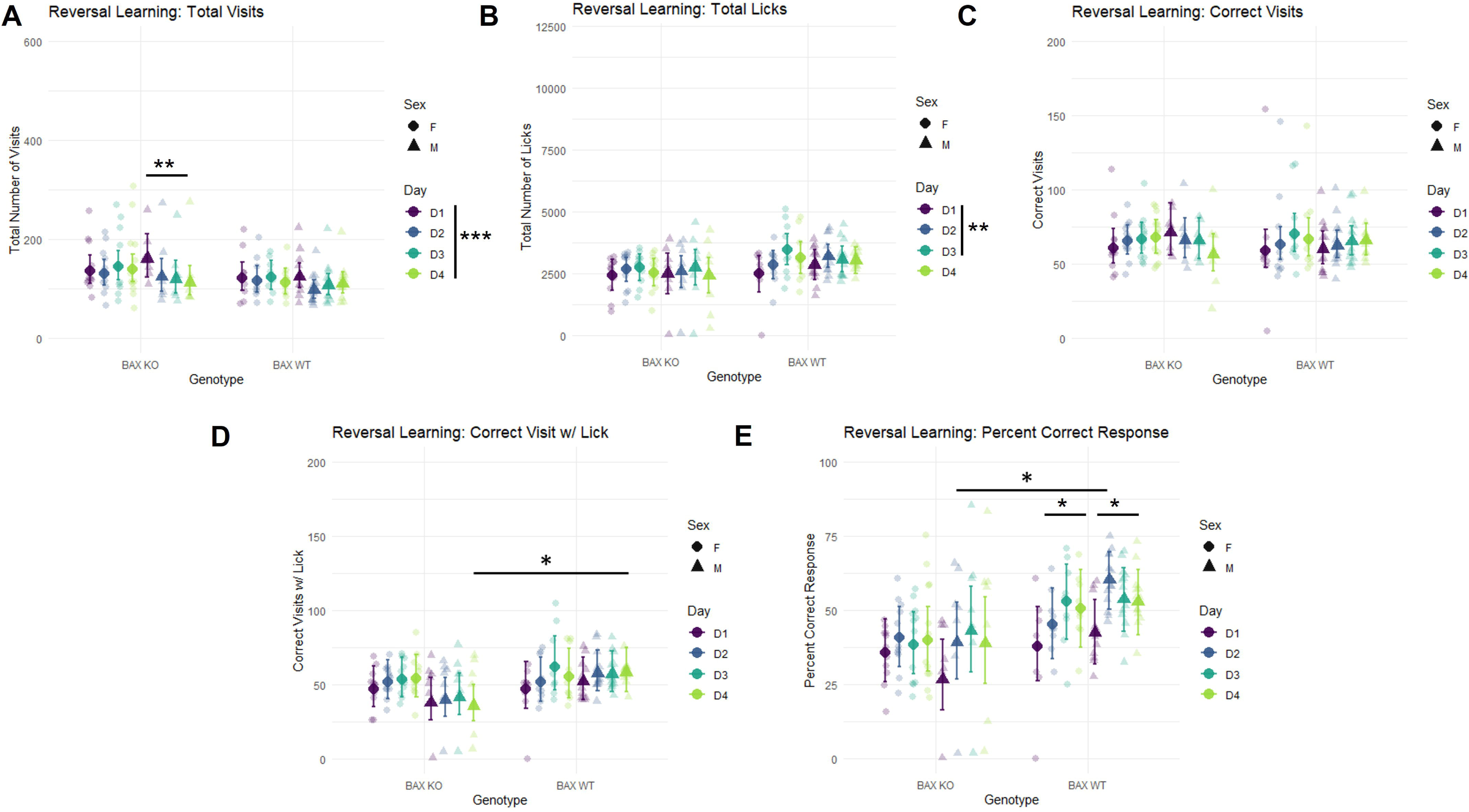
BAX WT mice performed better than BAX KO mice in reversal learning. (A) Total visits, (B) total licks, (C) correct visits, (D) correct visits with a lick, and (E) percentage of correct responses on days 1-4 (D1-D4). Small symbols, individual mice; large symbols, estimated marginal means (EMMs) with 95% confidence intervals (CIs). Circles: females; triangles: males. Vertical bars next to the keys mark main effects of day; horizontal bars mark pairwise comparisons. *p < 0.05, **p < 0.01, ***p < 0.001.

In terms of total licks (Fig. 6B), a significant effect of day was observed [F(3, 54) = 4.217; p = 0.009], on day 3 mice averaged 439.004 more licks than on day 1 (95% CI: [76.924, 801.084]; p = 0.010), while there was no significant difference on day 4 compared to day 1. Interestingly, all animals only showed a difference in licks from day 1 to day 3, but not from day 1 to day 4.

There were no significant effects on total correct visits (Fig. 6C).

There was a significant interaction between sex, genotype, and day on total correct visits with a lick [F(3, 53) = 3.031; p = 0.037] (Fig. 6D). On day 4, male BAX KO mice had 22.664 fewer correct visits with a lick compared to male BAX WT mice (95% CI: [-41.644, -3.684]; p = 0.020). On the same day, female BAX KO mice had 18.509 more correct visits with a lick than male BAX KO mice, however this difference was only near significance (95% CI: [-0.113, 37.130]; p = 0.051). There was a significant effect of day [F (3, 53) = 5.378; p = 0.003]. Across sex and genotype, on day 3 mice made 7.271 more correct visits with a lick (95% CI: [1.385, 13.157]: p = 0.008). However, there was no difference from day 4 to day 1. Male BAX KO mice made fewer visits with a lick than BAX WT mice.

As for percent correct response (Fig. 6E), there was a significant interaction between sex, genotype, and day [F (3, 44) = 5.332; p = 0.003]. Male BAX KO mice had a decrease of 21.2 percentage points in percent correct response on day 2 compared to male BAX WT mice (95% CI [-0.376, -0.048]; p = 0.013). On day 4, female BAX WT mice had an increase of 12.8 percentage points than day 1 (95% CI: [0.009, 0.247]; p = 0.029); male BAX WT mice had an increase of 10.4 percentage points than day 1 (95% CI: [0.008, 0.200]; p = 0.029). There was a significant effect of day on percent correct response [F (3, 44) = 15.443; p < 0.001]. There was an increase of 10.1 percentage point on percent correct response on day 4 compared to day 1 when looking across sex and genotype (95% CI [0.045, 0.156]; p < 0.001). There was an overall increase in percent correct responses indicating that all mice learned, however this effect was most prevalent in BAX WT mice as male and female BAX WT as they showed a significant difference from day 1 to day 4.

### Male BAX KO mice were better able to discriminate between shock and no-shock context compared to male BAX WT mice. BAX KO attenuated initial female low values

A significant interaction was found for sex, genotype, and day on discrimination score [F(4, 175) = 2.646; p = 0.035] (Fig. 7A). On day 1, female BAX KO mice scored 0.583 points higher than female BAX WT mice (95% CI: [0.183, 0.984]; p = 0.005). However, no significant differences were found between these groups on day 5. For male BAX KO mice, no significant differences were observed on day 1 compared to male BAX WT mice. However, by day 5, male BAX KO mice had an increase of 0.526 on discrimination score compared to male BAX WT mice (95% CI: [0.222, 0.831]; p = 0.001). Additionally, on day 5, male KO mice had an increase of 0.419 on discrimination score compared to female KO mice (95% CI [0.110, 0.728]; p = 0.001). On day 5 male BAX KO mice scored 0.776 points higher than day 1 (95% CI [0.153, 1.399]; p = 0.005. Likewise, on day 5 female BAX WT mice scored 0.876 points higher than day 1 (95% CI [0.337, 1.416]; p < 0.001. On day 2, female BAX KO mice had an increase of 0.594 points compared to day 1 (95% CI [0.077, 1.110]; p = 0.013). Similarly on day 2, female BAX WT an increase of 1.311 points compared to day 1 (95% CI [0.679, 1.943], p < 0.001).

**Figure 7.**
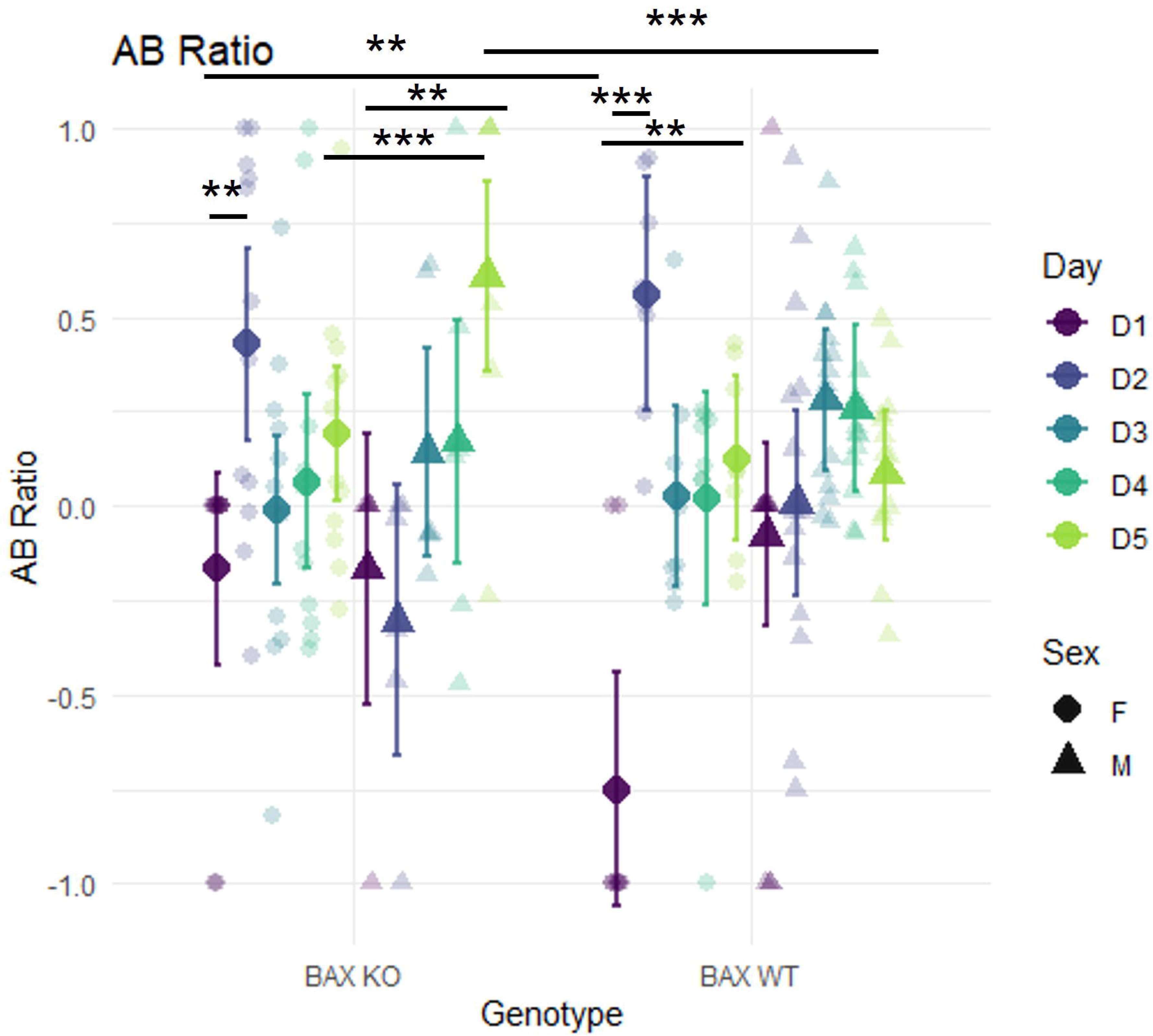
Male BAX KO mice discriminated between contexts better than male BAX WT mice in the A/B discrimination task. AB ratio on days 1-5 (D1-D5). Positive values indicate more freezing in the shock context; negative values indicate more freezing in the no-shock context. Small symbols show individual mice; large symbols show estimated marginal means (EMMs) with 95% confidence intervals (CIs). Circles: females; triangles: males. Horizontal bars mark pairwise comparisons. **p < 0.01, ***p < 0.001.

There was a significant interaction of sex and day [F (4, 175) = 7.229; p < 0.001]. On day 1, male scored 0.337 points higher than female mice (95% CI: [0.041, 0.632]; p = 0.026). On day 2, female mice scored 0.642 points higher than male mice (95% CI: [0.347, 0.937]; p < 0.001). There was a significant interaction effect of genotype and day [F (4, 175) = 3.035; p = 0.019). On day 5, across sex, BAX KO mice scored 0.295 points higher than BAX WT mice (95% CI: [0.088, 0.503]; p = 0.006). There was a significant effect of day [F (4, 175) = 9.176; p < 0.001]. On day 5, across sex and genotype, mice scored 0.542 points higher than day 1 (95% CI: [0.282, 0.802]; p < 0.001).

On the initial day female BAX KO mice performed better than female BAX WT mice, such that female BAX KO mice were freezing less in the no-shock than the shock context, achieving a more neutral response. On the second day female BAX KO and female BAX WT mice performed better than day 1, indicating a robust acute learning response. However, in female BAX KO mice, due to starting at a more neutral position, did not have a significantly different score from day 1 to day 5. Female BAX WT mice did show a difference due to their initial low score.

Male BAX KO mice performed better than female BAX KO mice on day 5. And they performed better than Male WT mice by day 5. They also performed better than female BAX KO mice on day 5. Male BAX KO mice performed better than day 1.

## Discussion

The goal of this study was to see if increasing adult hippocampal neurogenesis improves learning in male and female mice, and whether the benefit is the same in both sexes. We used BAX KO and BAX WT mice of both sexes and tested them in an Intellicage, which has the advantage that mice are tested in their home cage and are not handled by an experimenter. We first confirmed that BAX KO mice had more surviving newborn dentate granule cells than BAX WT mice, and this was true for both sexes. During the adaptation phases there were no differences between genotypes or sexes in the number of visits or licks the mice made. Differences only showed up during place learning, reversal learning, and the A/B context discrimination task. Male mice had a higher percent correct response during place learning, while female mice improved the most from day 1 to day 4. During reversal learning the BAX KO mice, especially the males, made fewer visits. In the A/B task, BAX KO improved discrimination only in male mice. These results partially support our hypothesis that increasing neurogenesis would benefit male mice more, but they also show that the effect depends on the task.

Studies that focus on neurogenesis often do not include female mice or look at sex-differences (Knudson, Hipkin and Snyder, 2022). As behavioral performance is influenced by testing environment and observer, the Intellicage system is a valuable tool. Although increasing neurogenesis has been associated with improved spatial learning and pattern discrimination, whether there are sex differences in this benefit remain unclear. In this study we used BAX KO and BAX WT mice of both sexes and ran them through a contextual spatial learning task in an Intellicage and a pattern discrimination task. We successfully increased neurogenesis by BAX KO; we did not see any sex differences. BAX KO mediated enhanced neurogenesis was not a surprise as this has been done before in a similar mouse model (Sahay *et al*., 2011; Besnard and Sahay, 2021). No sex differences have been found in baseline neurogenesis (Tsao *et al*., 2023). During the adaptation phase of the Intellicage, which included free adaptation (FA), door adaptation (DA), nosepoke adaptation (NPA) there were no difference between genotypes and sex on total visits and licks made. This is in line with what we saw in the open field test as there were no differences (Sup. Fig. 1). This also follows along with what was previously seen by Sahay et al. (2011), they did not see any genotype differences in BAX KO and BAX WT mice where both males and females were used. Furthermore, Tsao et al. (2023) did not see any sex differences in the open field test to indicate differences in activity. However, Mifflin et al. (Mifflin *et al*., 2021) who also used the Intellicage did see sex differences during the adaptation phase; male mice were making more visits and licks on the first day of adaptation although this was attenuated as adaptation continued. However, the strain of mice differed from the background strain of the mice used in this study (129/Svj versus C57BL/6J). When these two strains were directly compared 129/Svj showed less activity and a more anxious phenotype (Homanics, Quinlan and Firestone, 1999) which probably contributed to the sex differences that were seen by Mifflin et al. (2021). However, sex differences were not directly evaluated by Homanics et al. (1999). For our study, seeing no differences in adaptation meant that there were no baseline differences in Intellicage that had to be considered when evaluating place learning and reversal learning.

Place learning was conducted over the span of 4 days. Mice could only get water in a designated correct corner. During this phase, we saw that BAX KO mice were making more visits to all corners by day 4 than they were making on day 1, this was especially true for female BAX KO mice. As this difference was not present during the adaptation phase, this indicates that BAX KO mice were responding to the change in their environment, that is that water access was limited. Interestingly, Sahay et al. (2011) saw something similar. They only saw a genotype difference in the OFT when they included voluntary exercise. Although these two cases are not parallel, it shows that mice respond to changes in their environment. Overall, male mice were making more licks than female mice, however only female mice, BAX WT and BAX KO made a different number of licks from day 1 to day 4. Indicating that they were learning where to get water. Mifflin et al. (2021) also saw increase licks in male mice during place learning. However, unlike Mifflin et al. (2021), this increase was not present during adaptation. Thus, it is less likely that this overall increase in licks was due to a phenotype in male mice being thirstier. Interestingly, only male WT mice did not show an increase from day 1 to day 4 in total correct visits made.

However, this was probably due to a ceiling effect. Male mice had a higher percent correct response than female mice. Mifflin et al. (2021) also saw that male mice performed better at the place learning task than females. Our results go in line with meta-analysis that has shown that males perform better than female at spatial learning tasks (Yagi and Galea, 2019). However, this is not an instance where female mice did not learn. In fact, female mice showed the most improvement from d1 to day 4, in total licks and correct visits.

After place learning, mice were run through reversal learning where the opposite corner was chosen as the new correct corner. It is during this phase, where mice have to update where the correct corner is, is suggested to be most neurogenesis-dependent (Hernández-Mercado and Zepeda, 2022). Contrary to what we saw in place learning, mice were making fewer visits during reversal learning, with male BAX KO especially making less visits. There were no sex differences on number of licks made. Mice only made a significantly higher number of licks on day 3 than day 1, no difference on day 4. Following the pattern of decreased total visits, male BAX KO mice made less visits with a lick. The most interesting effect we saw was that there were no differences from day 1 to day 4 in BAX KO female and male mice, while there was a difference in BAX WT males and females. In fact, on day 2 male BAX KO mice performed worse than BAX WT mice on percent correct response. We did not see an improved performance on the initial day of reverse learning as seen by Barth et al. (2023) where mice with increased neurogenesis had an initial better performance that then attenuated. However, in a Morris water maze, which has been seen to be comparable to place learning paradigms in the Intellicage (Kiryk *et al*., 2020), Berdugo-Vega et al. (2020) saw that mice with a genotype for increased neurogenesis performed worse than WT mice on the first day of reversal learning. Mice did learn where they could get water as their licks increased. However, the decrease in total visits could be interpreted as a habituation of their environment, that is they no longer were curious as to what was happening in the cage corners.

Prior to placement in Intellicage, mice were run through a pattern discrimination learning task, contextual learning. In this task, mice are placed in two similar but distinct boxes, in one they get a shock and in the other they do not get a shock. The expectation is that mice learn to distinguish the no shock context and freeze less; and freeze more in the shock context. On day 5, Male BAX KO mice performed better than male BAX WT mice. This replicates the prior results that Sahay et al (2011) and Besnard & Sahay (2021) saw. On the initial day female BAX KO mice performed better than female BAX WT mice. This could be showing that increased neurogenesis could be attenuating the stressfulness of the new environment but there were no genotype and sex interaction in the TST (Sup. Fig. 2). There was an interesting effect were female BAX KO and BAX WT mice performed noticeably better on day 2 than day 1. Indicating something akin to single-trial learning in female mice. This could be due to a sensitivity to stress as indicated by female mice freezing more in the TST. However, this effect was not observed after day 2, which aligns with the prior literature showing that female mice are more prone to generalizing fear responses (Fleischer and Frick, 2023).

Our hypothesis that BAX KO would improve performance in male mice at spatial learning and pattern separation was only partially supported. BAX KO was only improved the performance of male mice in the pattern separation task. As for spatial learning, we saw a similar results as to what has been described in literature where male mice perform better at spatial learning tasks. However, this was only true in the place learning and not for reversal learning. The results from the reversal learning task leave an interesting question, why were mice making less visits. We believe that this may have something to do with the habituation or novelty response to the environment, and it could be tested directly using the novelty tasks that can be run in the Intellicage (Kiryk *et al*., 2020). Overall, our results show that increasing the survival of nDGCs does not improve learning in the same way for all mice. BAX KO helped male mice in the A/B context discrimination task, it increased the number of visits female mice made during place learning, and it was associated with fewer visits in male mice during reversal learning. This means that the effect of increased neurogenesis depends on the sex of the mouse and on what the task is asking the mouse to do. Future studies using these manipulations should include both sexes and should test more than one phase of learning.

## Acknowledgements

We thank the UNM Comprehensive Cancer Center and the Advanced Light Microscopy Shared Resource, NCI P30CA118100.

## References

Bankhead, P. et al. (2017) “QuPath: Open source software for digital pathology image analysis,” Scientific Reports, 7(1), p. 16878. Available at: 10.1038/s41598-017-17204-5.

Barth, K. et al. (2023) “EGFL7 loss correlates with increased VEGF-D expression, upregulating hippocampal adult neurogenesis and improving spatial learning and memory,” Cellular and Molecular Life Sciences: CMLS, 80(2), p. 54. Available at: 10.1007/s00018-023-04685-z.

Berdugo-Vega, G. et al. (2020) “Increasing neurogenesis refines hippocampal activity rejuvenating navigational learning strategies and contextual memory throughout life,” Nature Communications, 11(1), p. 135. Available at: 10.1038/s41467-019-14026-z.

Besnard, A. and Sahay, A. (2021) “Enhancing adult neurogenesis promotes contextual fear memory discrimination and activation of hippocampal-dorsolateral septal circuits,” Behavioural Brain Research, 399, p. 112917. Available at: 10.1016/j.bbr.2020.112917.

Fleischer, A.W. and Frick, K.M. (2023) “New perspectives on sex differences in learning and memory,” Trends in Endocrinology & Metabolism, 34(9), pp. 526–538. Available at: 10.1016/j.tem.2023.06.003.

Grieco, F. et al. (2021) “Measuring Behavior in the Home Cage: Study Design, Applications, Challenges, and Perspectives,” Frontiers in Behavioral Neuroscience, 15. Available at: 10.3389/fnbeh.2021.735387.

Gustus, K., et al. (2020) “Functional and Structural Correlates of Impaired Enrichment-Mediated Adult Hippocampal Neurogenesis in a Mouse Model of Prenatal Alcohol Exposure,” Brain Plasticity. Edited by K. Nixon, 6(1), pp. 67–82. Available at: 10.3233/BPL-200112.

Hamson, D.K., Roes, M.M. and Galea, L.A.M. (2016) “Sex Hormones and Cognition: Neuroendocrine Influences on Memory and Learning,” Comprehensive Physiology, 6(3), pp. 1295–1337. Available at: 10.1002/j.2040-4603.2016.tb00710.x.

Hernández-Mercado, K. and Zepeda, A. (2022) “Morris Water Maze and Contextual Fear Conditioning Tasks to Evaluate Cognitive Functions Associated With Adult Hippocampal Neurogenesis,” Frontiers in Neuroscience, 15. Available at: 10.3389/fnins.2021.782947.

Homanics, G.E., Quinlan, J.J. and Firestone, L.L. (1999) “Pharmacologic and Behavioral Responses of Inbred C57BL/6J and Strain 129/SvJ Mouse Lines,” Pharmacology Biochemistry and Behavior, 63(1), pp. 21–26. Available at: 10.1016/S0091-3057(98)00232-9.

Jonasson, Z. (2005) “Meta-analysis of sex differences in rodent models of learning and memory: a review of behavioral and biological data,” Neuroscience & Biobehavioral Reviews, 28(8), pp. 811–825. Available at: 10.1016/j.neubiorev.2004.10.006.

Judd, J.M. et al. (2024) “Modifying reaction time tasks parameters in the automated IntelliCage identifies heightened impulsivity and impaired attention in the 3xTg-AD model of Alzheimer’s disease,” Frontiers in Aging Neuroscience, 16, p. 1466415. Available at: 10.3389/fnagi.2024.1466415.

Kas, M.J.H. and Van Ree, J.M. (2004) “Dissecting complex behaviours in the post-genomic era,” Trends in Neurosciences, 27(7), pp. 366–369. Available at: 10.1016/j.tins.2004.04.011.

Kiryk, A. et al. (2020) “IntelliCage as a tool for measuring mouse behavior - 20 years perspective,” Behavioural Brain Research, 388, p. 112620. Available at: 10.1016/j.bbr.2020.112620.

Knudson, C., Hipkin, S. and Snyder, J. (2022) *Adult neurogenesis studies primarily use males*. figshare. Available at: 10.6084/M9.FIGSHARE.19319849.

Kyriakou, E.I. et al. (2018) “Home-cage anxiety levels in a transgenic rat model for Spinocerebellar ataxia type 17 measured by an approach-avoidance task: The light spot test,” Journal of Neuroscience Methods, 300, pp. 48–58. Available at: 10.1016/j.jneumeth.2017.08.012.

Mifflin, M.A. et al. (2021) “Sex differences in the IntelliCage and the Morris water maze in the APP/PS1 mouse model of amyloidosis,” Neurobiology of Aging, 101, pp. 130–140. Available at: 10.1016/j.neurobiolaging.2021.01.018.

Rodriguez, A.M. et al. (2026) “Fluoxetine for Prenatal Alcohol-Exposed Mice: Addressing Impaired Enrichment-Mediated Neurogenesis.” Neuroscience. Available at: 10.64898/2026.03.13.711456.

Sahay, A. et al. (2011) “Increasing adult hippocampal neurogenesis is sufficient to improve pattern separation,” Nature, 472(7344), pp. 466–470. Available at: 10.1038/nature09817.

Sousa, N., Almeida, O.F.X. and Wotjak, C.T. (2006) “A hitchhiker’s guide to behavioral analysis in laboratory rodents,” *Genes*, Brain and Behavior, 5(s2), pp. 5–24. Available at: 10.1111/j.1601-183X.2006.00228.x.

Tsao, C.-H. et al. (2023) “The influence of sex difference on behavior and adult hippocampal neurogenesis in C57BL/6 mice,” Scientific Reports, 13(1), p. 17297. Available at: 10.1038/s41598-023-44360-8.

Weigard, A., Loviska, A.M. and Beltz, A.M. (2021) “Little evidence for sex or ovarian hormone influences on affective variability,” Scientific Reports, 11(1), p. 20925. Available at: 10.1038/s41598-021-00143-7.

Yagi, S. and Galea, L.A.M. (2019) “Sex differences in hippocampal cognition and neurogenesis,” Neuropsychopharmacology, 44(1), pp. 200–213. Available at: 10.1038/s41386-018-0208-4.

