## Supplemental Figures for "Sex differences in the effects of enhanced adult hippocampal neurogenesis on pattern discrimination and Intellicage spatial learning in Bax-knockout mice"

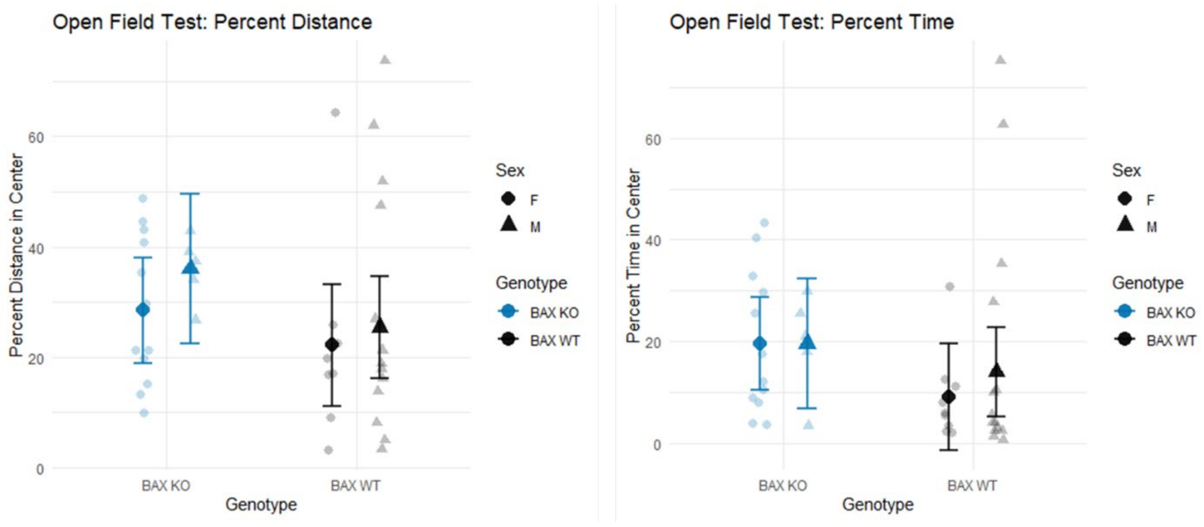


**Sup. Figure 1. Center exploration in the open field test did not differ by genotype or sex.** (A) Percentage of total distance traveled in the center and (B) percentage of time spent in the center of the open field. Small symbols show individual mice; large symbols show estimated marginal means (EMMs) with 95% confidence intervals (CIs). Circles: females; triangles: males. BAX KO: blue; BAX WT: black.

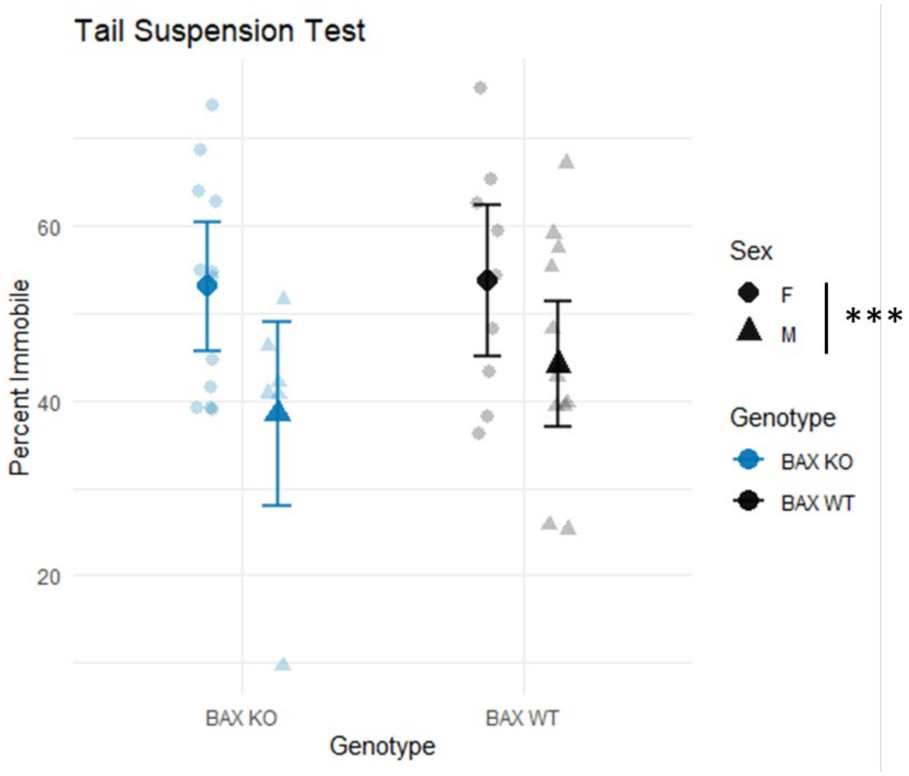


**Sup. Figure 2. Female mice spent more time immobile than male mice in the tail suspension test.** Percentage of time spent immobile. Small symbols show individual mice; large symbols show estimated marginal means (EMMs) with 95% confidence intervals (CIs). Circles: females; triangles: males. BAX KO: blue; BAX WT: black.***$p < 0.001$, main effect of sex.
